# In Search of the Perfect Model: How Cancer Cell Lines Relate to Native Cancers

**DOI:** 10.1101/2024.05.15.594310

**Authors:** Rahel Paloots, Ziying Yang, Michael Baudis

## Abstract

Cancer cell lines are frequently used in biological and translational research to study cellular mechanisms and explore treatment options. However, cancer cell lines may display mutational profiles divergent from native cancers or may be misidentified or contaminated. We explored how similar cancer cell lines are to native cancers to find the most suitable representations for the corresponding diseases by utilising large collections of copy number variation (CNV) profiles and applied machine learning (ML) algorithms to predict cell line classifications.

Our results confirm that cancer cell lines indeed accumulate more mutations compared to native cancers but retain similar CNV profiles. We demonstrate that many relevant oncogenes and tumor suppressor genes are altered by CNV events in both cancers and their corresponding cell lines. Based on the similarities between the two groups and the predictions of the ML model, we provide some recommendations about cell lines with good potential to represent selected cancer types in *in vitro* studies.

## Introduction

Derived from cellular samples of primary or recurring tumors, hematopoietic neoplasias, or cancer metastases, cancer cell lines serve as tractable *in vitro* representations of human malignancies, enabling researchers to dissect cancer biology and evaluate potential therapeutic interventions. They are popular models due to established handling procedures and relatively low cost. To establish a cancer cell line, cells are isolated from a native cancer *e*.*g*. a solid tumor or bone marrow in case of some lymphoid neoplasms. The ability of many cancer cells to divide perpetually is exploited for propagating them indefinitely for recurrent use in studies, potentially over the course of decades. For example, the first cancer cell line ever established, HeLa (1951) (1), is still one of the most commonly used *in vitro* model systems.

Since the establishment of HeLa and besides individual cell lines established in research projects and maintained at individual institutions, thousands of cell lines have been made available through commercial services, promising precise matching of cell lines to disease types. Many fixed sets of cell lines have been created to model different cancer types in a consistent manner under experimental conditions, e.g. for comparing drug activity or effects of targeted genetic modifications. A widely used collection of cancer cell lines is the NCI-60 panel that represents 9 distinct cancer types (2).

Cancer tissues are complex and genetically heterogeneous, frequently comprising various clonal populations (3, 4). Genomic aberrations in cancers enable disease progression as well as metastatic growth. A major class of these genomic alterations are copy number variations (CNVs) where large regions in the genomes have been modified by amplification or deletion of a section. These CNV profiles are also cancer type specific, *e*.*g*. duplication of chromosome 13 in colorectal carcinomas (5– Moreover, genomic alterations in cancers depend on the grade as well as the stage of the disease (3), suggesting different subtypes can be detected within disease classification. For example, medulloblastomas have four well known and characterized subtypes. Two of these well-known subsets are based on the expression of Wnt and Shh genes, the other two groups are “group 3” and “group 4” since the biology behind these types is not clear (8).

Due to their restricted origin, cancer cell lines inherently capture only a limited subpopulation of the original tumor’s genetic diversity. This inherent limitation is further compounded by the selective pressure exerted by in vitro culture conditions. *In vitro* systems also lack the crucial interactions with other cell types that help shape neoplastic growths. Therefore, under these disparate conditions, cancer cell lines obtain novel features, such as bear a larger amount of copy number alterations compared to primary cancers (9, 10).

In this study, we evaluated cancer cell lines of different types and compared their profiles to their respective native cancer types. We applied statistical and machine learning models to identify CN patterns within 31 distinct diagnostic terms. We give an overview of the genomic differences between cancers and their cell lines and unravel the key differences in the genomic features of the two groups. Based on these results, we suggest the best available cell lines per diagnostic group.

## Methods

### Input Data

Cancer cell line CNV data used in this study originates from cancercelllines.org our knowledge resource for cancer cell line variants. This database currently includes about 5,600 individual CNV samples (11). Progenetix, our source for native cancer samples includes around 112,000 cancer related CNV profiles (12). Following an initial analysis of cell line samples, a subset of 31 cancer types was chosen for further investigation. This selection was guided by the distribution of representative samples within the cancer cell lines set.

To reduce the impact of samples with limited quality of their CNV profiling data, all cell line samples were assessed visually and hlproblematic samples were flagged. Native cancer samples without any detected segments were excluded from the dataset automatically. Both native cancer and cell line samples labeled as Unspecified Tissue (NCIT:C132256) were also excluded from the dataset. A set of NCIT cancer types with a sufficient number of cancer cell line samples was used in the similarity assessment and machine learning models 2.

### Binning of CNV Call Values

To bring cell line and tumor data to a uniform format, CNV data was transformed into a binned matrix. All bins are of equal specified length and represent an area in the genome. Inside each bin, CNV coverages are calculated. Duplication and deletion coverages of the bins are calculated separately. All CNV gains or losses in a bin are counted and the fraction of CNVs covering the bin are calculated. All bins with CNV gain fractions are then saved into an array, followed by the bins with deletion coverage fractions. An open-source Python package (bycon) was used for bin calculations. Genomic intervals were then calculated for different bin sizes from 1 to 10Mb. A bin size of 5 Mb was selected for further evaluations to avoid biases due to small and very large segments in the genome. This resulted in 635 bins for duplications and 635 bins for deletions, total 1270 bins.

Based on genomic bins the bycon package also supports the creation of CNV frequency maps, representing the occurrence of a CNV (gain or loss) for each segment as a percent value for all selected samples (%). For example, if all lung carcinoma samples have a duplication in 8q, the frequency in this region would be 100%. CNV frequencies are calculated based on a 1 Mb bin size to capture locations of focal events. Calculated frequencies were then used for similarity assessments and visualizations.

### CNV Coverage Calculation

CNV coverage of cell line and tumor samples were retrieved from progenetix and cancercelllines databases. CNV coverage fraction is the amount of the genome that is affected by structural variants (gains and losses). To assess the levels of structural variants between cell lines and tumors of the same cancer types, the average CNV coverage fractions and subsequently fold changes between the two groups were calculated. Pre-calculated CNV coverages for both groups are included in cancercelllines and progenetix databases.

### Cosine Similarity

Cosine similarity is a suitable measure for bins with values between 0 and 1 due to its scale invariance, which ensures consistent comparison regardless of vector magnitude. Its angle-based approach captures directional information, making it effective for assessing relative proportions or relationships between values, particularly in sparse datasets. Additionally, its normalized output provides easy interpretation and comparison across different datasets or dimensions. Pairwise cosine similarities between samples were calculated to detect outliers and assess the similarity between instances of the same cell line. Additionally, similarities between different cell lines as well as cell lines and native cancers were determined. Cosine similarity was calculated by using open source python package scikit-learn (Version: 1.2.2). To assess the overall similarity of cell line and native cancer profiles, cosine similarities of CNV frequencies were calculated.

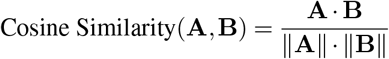

### Supervised Machine Learning Algorithms

We trained support vector machine (SVM) and random forest (RF) algorithms on native cancer samples to be able to predict cancer cell line diagnostic classifications based on the profiles of neoplasms. Python package scikit-learn (Version: 1.2.2) SVM classifier with rbf kernel and RF classifiers (n_estimators=200) were used for training and prediction. Data matrices with bin sizes from 1-10 Mb were tested and 5 Mb bin size selected for further use. To find the accuracy of differentiating between cancer cell line and native cancer, equal numbers of neoplasia instances to cancer cell lines were picked at random, as the number of neoplasia samples is greater. To predict cell line diagnostic labels based on tumor profiles, machine learning models were trained on selected cancer types. Cancer types were chosen based on the number of cell line samples available per NCIT classification term and child terms of the types were included (Table 2). Testing size for all models was 0.25 with random_state=10. Table **??** outlines the individual stages of various models.

### Feature Importances

We utilized scikit-learn’s PermutationImportance to determine the contributions of each feature to predictions. The script fits a PermutationImportance object to the dataset and computes feature importances using permutation tests. Class feature importances (“Cell Line” and “Tumor”) were then calculated for both SVM and RF models. Feature importance results included training with “Cell Line”-“Tumor” of equal numbers to identify general differences between all neoplasias and cell lines. We identified features important for distinct cancer types by training the algorithms on all cell line and neoplasia samples of the same diagnosis. The feature importances of RF model were used for this analysis because of the inherent feature importance measures of the model and the ability to capture complex relationships. Identified features were then mapped to the COSMIC set of signature genes https://cancer.sanger.ac.uk/signatures/downloads/ (last accessed 2023-08-28) to identify relevant genes in these bins.

### Matching Cell Lines with Cancer Subtypes

We partitioned cancer data by using the K-means clustering algorithm. Each cancer type (primary cancer samples only) was partitioned into clusters using 2-20 number of clusters. Median sample of each cluster in the partitioned data was calculated and the median sample of each cell line as well to represent each group. Then, cosine similarity between the medians was calculated. Only similarities equal to and above 0.7 were analyzed further.

### Plotting and Visualization

All frequency maps and CNV profile plots were created with progenetix/cancercelllines online software tools and bycon package software. Other graphs were created with python packages plotly, seaborn and matplotlib.

## Results

### Cancer cell lines display limited heterogeneity

A cancer cell line derived from a single source would exhibit minimal genetic variation within its population and within the samples of the same cell line. Even though cancer cell lines are not necessarily monoclonal, they still only represent a small subset of a tumor cell population. To determine the uniformity of the samples of the same cell line, we performed pairwise similarity calculations for all cell lines with at least 3 samples. Then, we compared computed indices to CNV sample plots to ascertain the efficiency of the measure by visual assessment. Figure 1 depicts 2 analyzed cell lines: prostate small cell carcinoma cell line NCI-H660 (CVCL_1576) and lung adenosquamous carcinoma cell line NCI-H596 (CVCL_1571). NCI-H660 is an example of a largely homogenous cell line with all similarity scores above 0.7 and there is a visual congruence among the samples on the CNV plot as well. NCI-H596 on the other hand portrays 2 distinct subsets that also brought about lower scores. Both subsets exhibit high similarity within the group but diverge from each other significantly. These results suggest that our measure can accurately determine the similarity of individual samples.

**Fig. 1.**
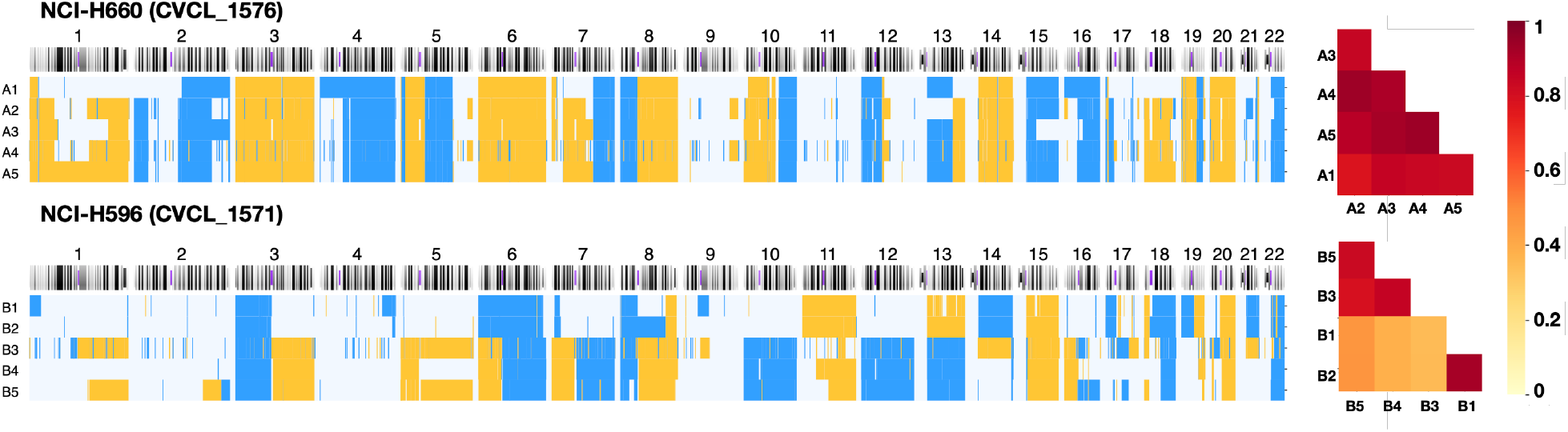
CNV sample plots and similarity heatmaps for cell lines NCI-H660 (top) and NCI-H598 (bottom), indicating the regional coverage by copy number gains (yellow) and losses (blue) for chromosomes 1-22. For each of the 2 cell lines 5 individual instances are shown to visualize similarities and differences in CNV events. Inter-sample CNV cosine similarities are displayed on the right.

To disclose the homogeneity of the cell lines of a distinct cancer type, we calculated pairwise similarities of all samples within the same cell line (at least 3 samples) and cancer type (Supplementary Fig. A1). We demonstrate a quantifiable level of homogeneity in cell line samples for the majority of cancer types. The average level of homogeneity across these samples surpasses 0.6. Lower similarity was detected for cervical carcinoma and fibrosarcoma cell lines. These results suggest that samples of the same cell line are overall relatively homogeneous. To detect any outliers, we set the similarity threshold within the same cell line to 0.6. This threshold was set based on visual assessment of single sample plots in combination with calculated similarity indices. Therefore cell lines with at least one paired score below 0.6, would have an outlier. 53% of the cell lines had at least one outlier and only 2% diverged greatly with no similarities above threshold. Overall, the data indicated limited variability between the instances of the same cell line.

### Cancer cell lines have higher CNV coverage compared to tumors of the same cancer type

Studies in breast and ovarian carcinoma cell lines have established that compared to tumors, cell lines accumulate more mutations (9, 10). For a systematic assessment of how cancer cell line genomes relate to neoplasias, we compared the CNV coverage fractions of both genomes. As expected, the CNV coverage in most cancer types was higher in cancer cell lines than in neoplasias (Fig.2) with the average fold change of 2.6. The exception was fibrosarcoma where fold change was below 1 and CNV coverage in tumors was greater. In chronic myelogenous leukemia (CML) with BCR-ABL a 15-fold CNV fraction change was detected. This could be due to most cell lines being in the “blast phase” of CML when they acquire a higher mutation load due to increased genomic instability (13). The link between increased mutation load and blast phase becomes evident when considering the source of CML cell lines used in research, *e*.*g*. a study by Drexler et al. (2000) where most analyzed CML cell lines originated from patients in blast crisis (14). Interestingly, all three highest fold change values are in different types of leukemia, highlighting the importance of genetic fusion events in many hematopoietic leukemias. Another important fusion in leukemias is PML-RARA in acute myeloid leukemia (AML) but a variety of genes have been reported to be involved in gene fusions in leukemias (15).

### Native cancers are highly heterogeneous but exhibit similar CNV profiles to cancer cell lines

It has been reported that despite harboring a higher burden of CNVs compared to their tissues of origin cancer cell lines remarkably retain a similar CNV signature to native cancers (9, 10). To systematically assess the concordance between neoplasia and cell line samples, we calculated pairwise similarity scores based on CNV coverage. Our analysis of the comparisons between cell lines and neoplasias, along with the internal consistency of the neoplasia data set, revealed scores that were lower than what we had anticipated.(Fig.3). Heterogeneity was higher within neoplasia samples than in instances of different cell lines of the same cancer type (Fig. 3, Supplementary Fig. A3). Standard deviations of similarities were higher for neoplasias in all cancer types except acute myeloid leukemia.

Pairwise comparisons of samples illustrate the heterogeneity but fail to reveal recurring patterns of similarity. Therefore, we compared the CNV frequency maps of neoplasias and corresponding cell lines. Frequency maps represent the occurrence of CNVs in the dataset in percentages indicating the presence of CNVs in the proportion of samples. We calculated a similarity index based on CNV frequency data of cell lines and neoplasias and found high similarity indices for most cancer types (Supplementary Table A2). Clearly detectable patterns emerge between cell lines and its native cancer upon visual inspection of frequency plots as well (Supplementary Fig. A2). Our results confirm that both breast carcinomas and ovarian carcinomas exhibit similar CNV patterns to their cell lines (9, 10). Even though melanoma has bee reported as a highly heterogeneous disease (16, 17), it yielded a high similarity score (Supplementary Table A2, Supplementary Fig. A2) indicating an overall good representation of genomic patterns in aggregated cell line data.

### Emerging cancer CNV patterns can be used to determine the origin of some cancer cell lines

Given the high occurrence of characteristic CNV patterns in a cancer type, our goal was to employ a method to forecast the diagnostic classification of a cell line. For that we trained a support vector machine (SVM) model on the 31 cancer types (Supplementary Table A1). In the evaluation of the model’s performance we found that for some diagnostic groups the CNV based diagnostic prediction was overall successful with the best results observed for breast adenocarcinoma, glioblastoma and colorectal carcinoma (Supplementary Table A3).

We then applied the trained model to predict the diagnostic classifications of our collection of cell line CNV profiles. The prediction accuracies of the cell lines were largely similar to testing - highest percentage of correctly predicted samples belonging to breast adenocarcinoma, glioblastoma and colorectal carcinoma. Interestingly, the percentage of correctly predicted cell lines is higher for breast adenocarcinoma cell lines than tumor-tumor predictions, indicating that these cell lines are good representations of the majority of the disease. On the contrary, while the testing accuracy was the highest for acute myeloid leukemia (AML) (90.51%), the prediction accuracy of AML cell lines was below 20%. In fact, the results for all leukemia types in our data sets revealed poor performance. The fold change of CNV coverage was also the highest among leukemias (Fig. 2), suggesting poor representation of the majority of native leukemia samples.

**Fig. 2.**
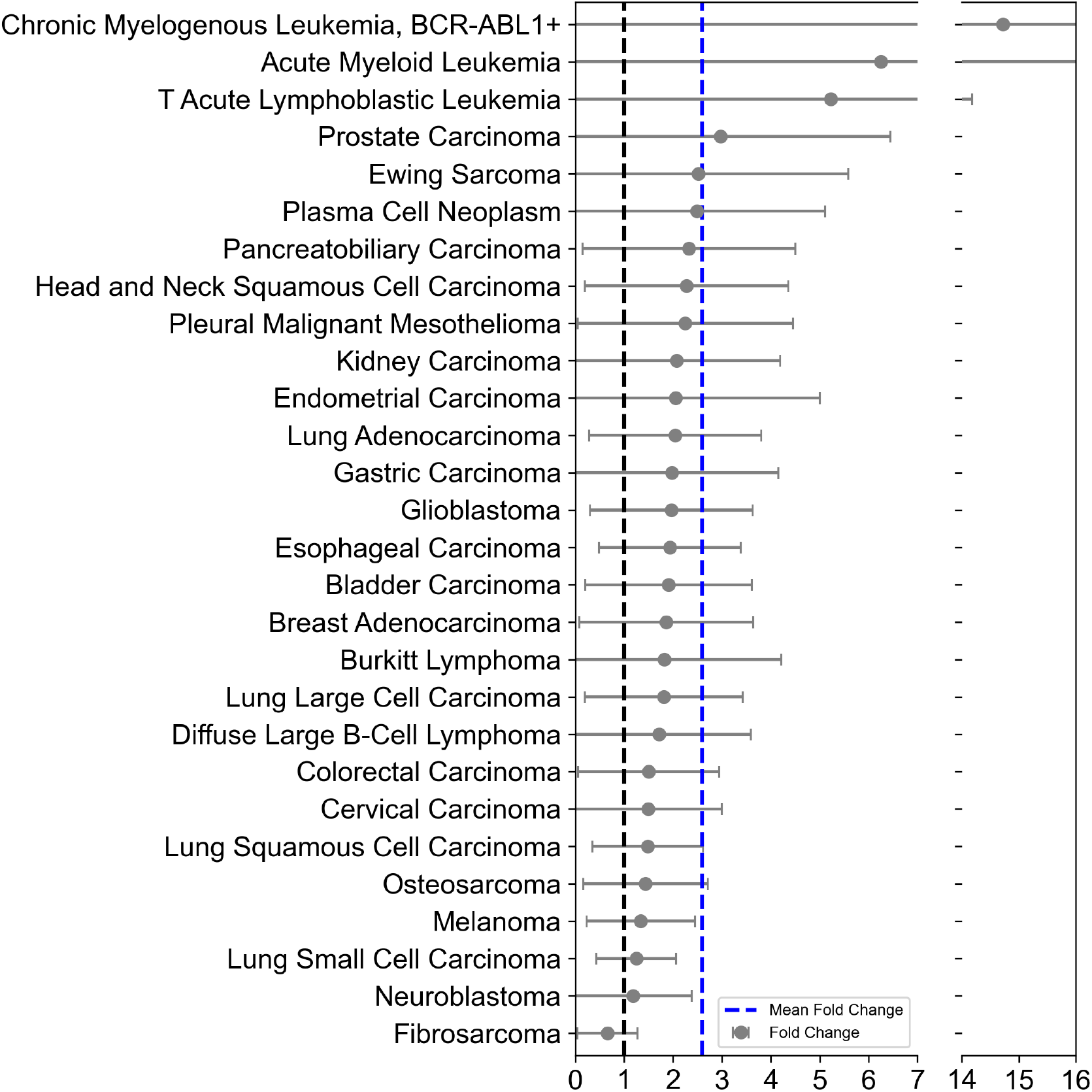
Comparison of CNV coverage fractions between cell lines and patient derived samples from the same diagnostic groups. Fold change calculated per cancer type: average CNV coverage of cell line samples/average CNV coverage of native cancer samples. Error bars represent standard errors.

**Fig. 3.**
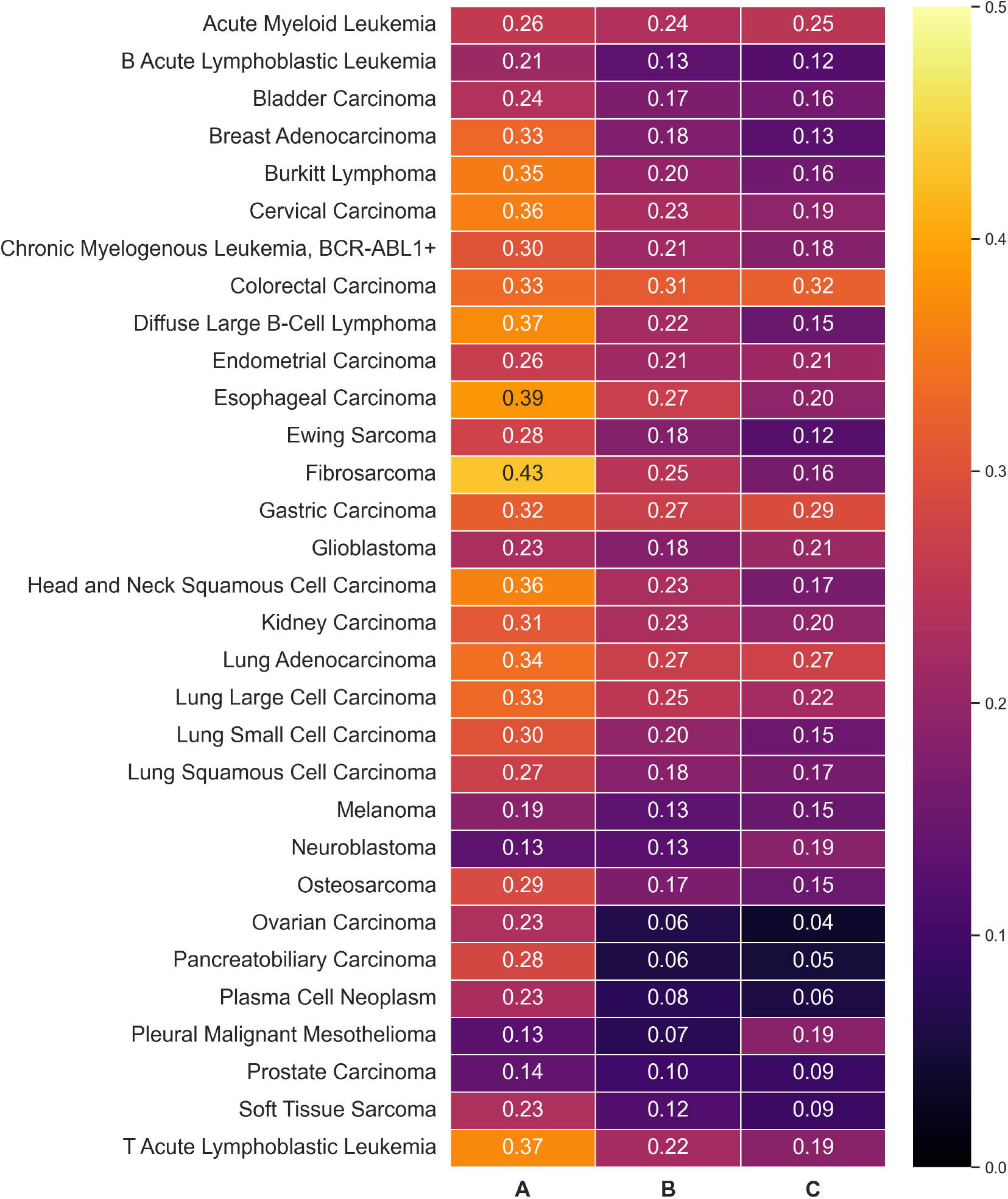
Similarity heatmap of comparisons between cell lines and neoplasias. Each column shows average pairwise similarity for the cancer type: A - samples of different cell lines, B - cell line samples vs neoplasia samples, C - neoplasia samples.

### Co-occurrence of relevant cancer genes in features important for class determination

To interpret the performance of our machine learning model, we employed a feature importance analysis upon its implementation. This analysis identified the most significant regions within the genome that contribute to the model’s classification capabilities.

Our initial objective was to examine whether any genomic features could be identified as influential in classifying “neoplasia” or “cell line”. Therefore, we collected all cell line samples in our dataset and randomly picked the same number of neoplasia samples. This step ensured the same number of samples to avoid introducing bias to our model. Distinguishing “neoplasia” from “cell line” with 89% accuracy (testing), this model also highlighted the most relevant features in the process (Fig. 4) which curiously include relevant cancer genes with functional involvement in many cancer types, such as TP53.

**Fig. 4.**
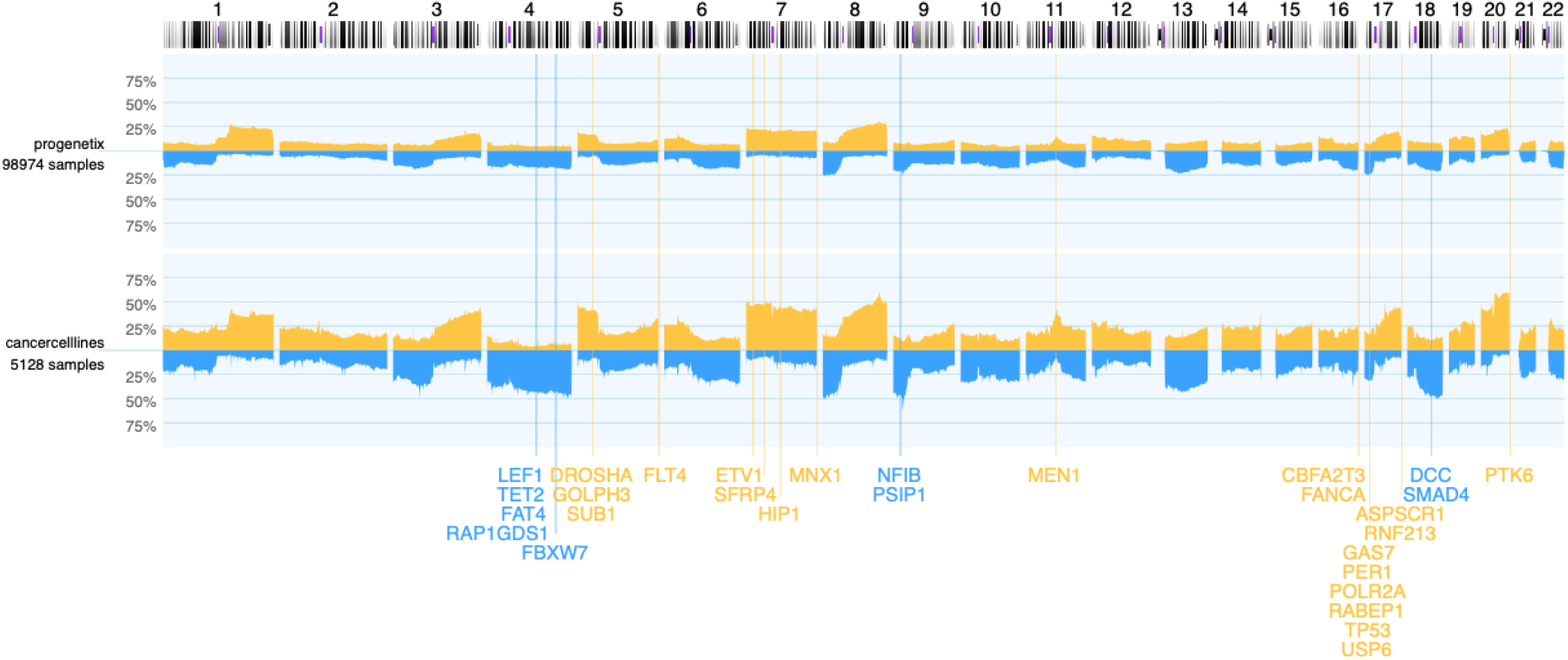
Frequency maps of all neoplasias and cancer cell lines. Blue - deletions, yellow - copy number gains, as percent of occurrence in analyzed samples. Locations of the top 30 genes with differences between cell lines and primary tumors in the ML model are indicated.

Next, we analyzed the important features in our selected 31 cancer types, by training our models with both cell line and neoplasia samples and finding the features relevant for each cancer type. Then, we matched identified features (genomic bins) with COSMIC cancer gene set, to identify any underlying oncogenes or tumor suppressor genes in the region. Of all the identified features, 56% are duplications and 44% are deletions. We then sorted the features by including only features with highest average values and that exist in at least 4 cancer types to have an overview of shared relevant features in the cancer types (Fig. 5). 42 out of 50 top shared features are duplications, indicating that duplications carry more weight in class identification. Many of these duplicated features include important genes such as BRCA1, EGFR, MAPK1. Wildtype-BRCA1 is a tumor suppressor gene and mutated BRCA1 increases risk for breast and ovarian carcinoma (18). BRCA1 is detected as an important feature for breast adenocarcinoma but not ovarian carcinoma (Fig. 5). EGFR-epidermal growth factor receptor, is an oncogene that promotes tumor progression. Notably, EGFR over-expression has been detected in lung adenocarcinomas but not in small cell lung carcinomas (SCLC) (19). Duplications in EGFR in lung adenocarcinomas and SCLCs are not marked as important features in our dataset. However, EGFR duplications are important features for determining breast adenocarcinoma sample types and breast cancers are also known to express EGFR (Fig. 5) (19). Another potent oncogene in cancers, particularly bladder carcinomas is FGFR3 (20). Interestingly, FGFR3 deletions are highlighted as important features for class detection in 5 cancer types, including bladder carcinomas (Fig. 5).

**Fig. 5.**
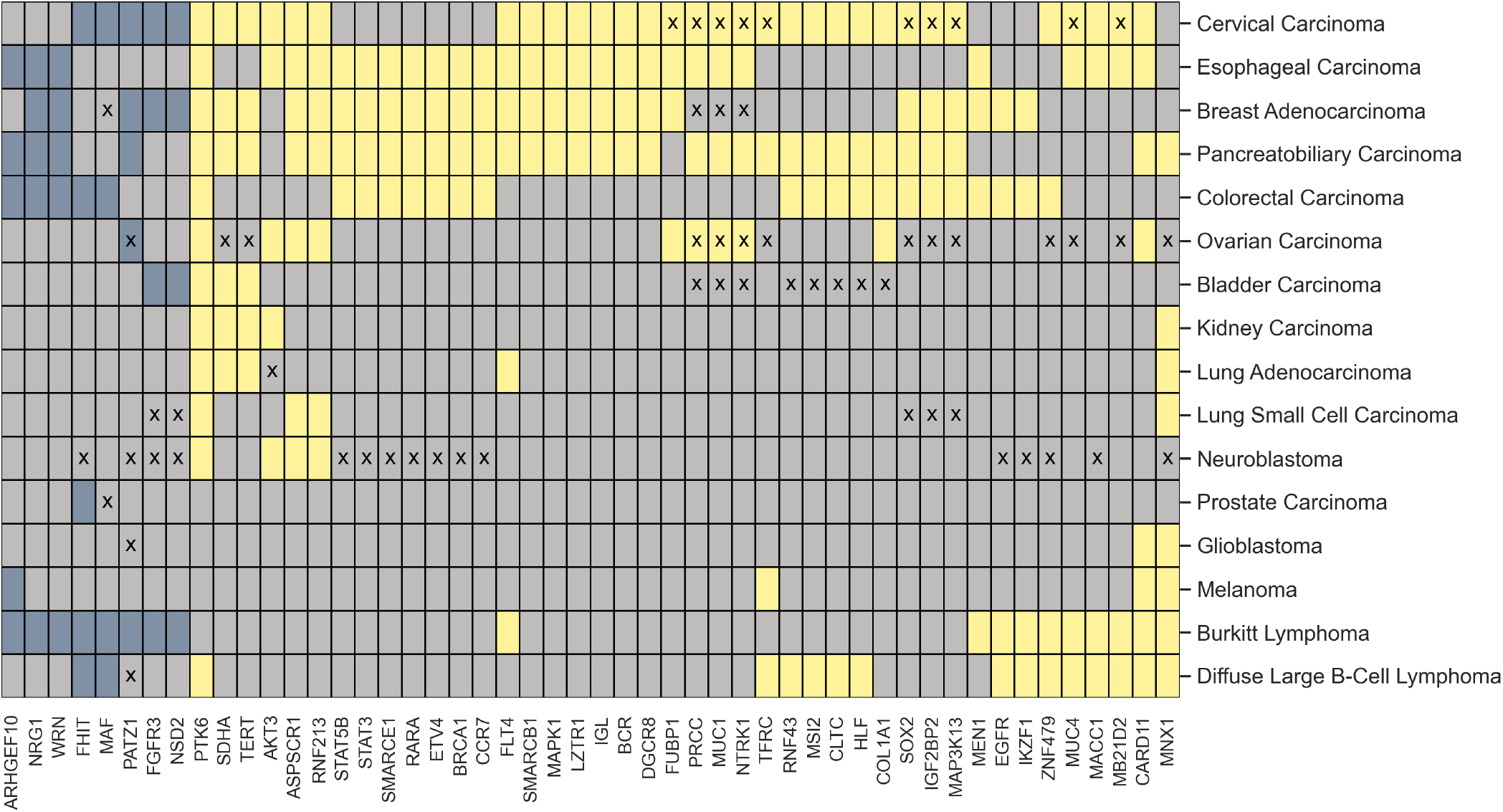
Known oncogenes and tumor suppressor genes important for class determination. Deleted genes are marked with blue and amplified genes with yellow, grey indicating no change in copy numbers. Genes exhibiting a higher incidence in native cancers relative to cell lines are denoted by ‘x’ on the heatmap.

Overall, the frequency of these features is lower in native cancers than in cell lines. This aligns with the known higher mutation rate observed in native cancers (Fig. 10, Fig. 11). While the frequencies of other features varied, the occurrence of mutations in certain genes remained consistent, e.g. COL1A1 in neuroblastomas and FHIT in lung small cell carcinomas. However, these genes were not deemed relevant for the classification algorithm.Different copy numbers in native cancers and cancer cell lines can be attributed to cancer cell lines only representing a small sub-population of the disease. The frequent use of cell lines derived from the same individual can lead to genetic drift, exacerbating these differences. Additionally, changes in copy numbers are observed in*in vitro* environments.

Our results suggest that several relevant cancer genes are an integral part for the determination of the class in our model. These genes and features are present in both neoplasias and cancer cell lines but at significantly higher levels in cell lines. These high level CN changes in cancer cell lines make them excellent models for studying the effects of these genes and testing for possible pharmaceuticals.

### Modeling Tumor Heterogeneity: The Utility of Cell Lines

Cancers are heterogeneous diseases that consist of multiple subsets. For example, medulloblastoma is a form of brain cancer that includes several distinct subtypes with characteristic mutational profiles (8). Another well-known heterogeneous cancer type is melanoma (17) where a large population of subclones have been detected (21). Based on known intratumoral heterogeneity, we hypothesized that certain cell lines would more accurately represent the unique molecular characteristics of specific cancer subsets. Therefore, we partitioned our neoplasia data using K-means clustering and picking the median sample of each cluster. We then matched these cluster medians to median samples of cell lines. Selecting the median sample allowed us to establish an “average” representation for each group. Indeed, we were able to match some subsets to some cell lines with high similarity, including melanomas and lung small cell carcinomas. For instance, Figure 6 displays a subset of lung small cell carcinoma tumor and cell line samples. We were able to identify three distinct cell lines that would be the best representations of this tumor subpopulation. Additionally, the subsets of melanoma, colorectal carcinoma, glioblastoma and kidney carcinoma matched with high similarity to several cell lines (Supplementary material).

**Fig. 6.**
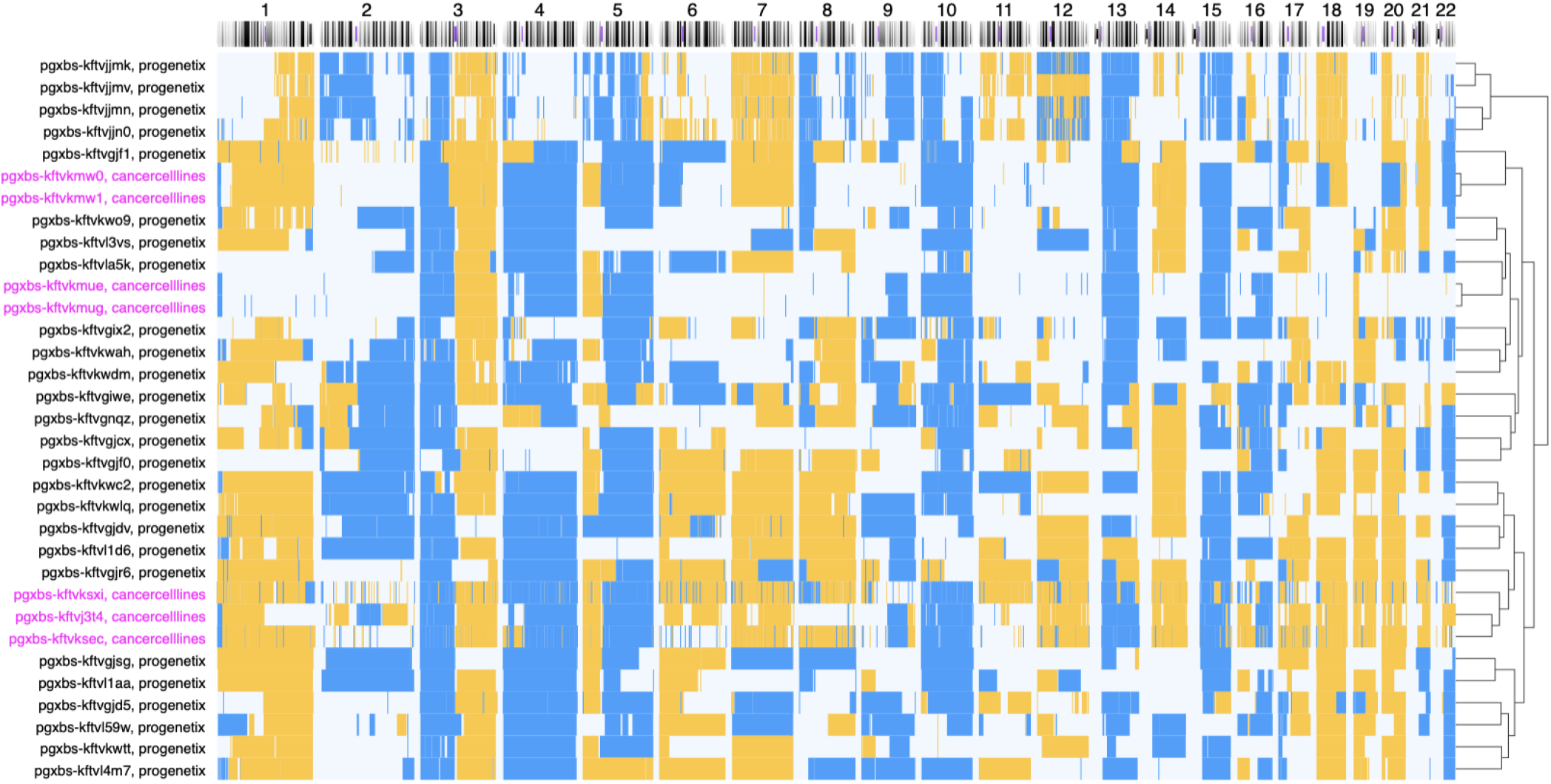
Clustered sample plots of similar cell line and neoplasia subset samples. Here, a subset of small cell lung carcinoma (SCLC) samples is interspersed by instances of SCLC cell in the unsupervised clustering analysis, indicating the high degree of CNV pattern similarity between some SCLC samples and those in vitro models. Cell line samples are indicated in pink and neoplasia samples in black.

**Fig. 7.**
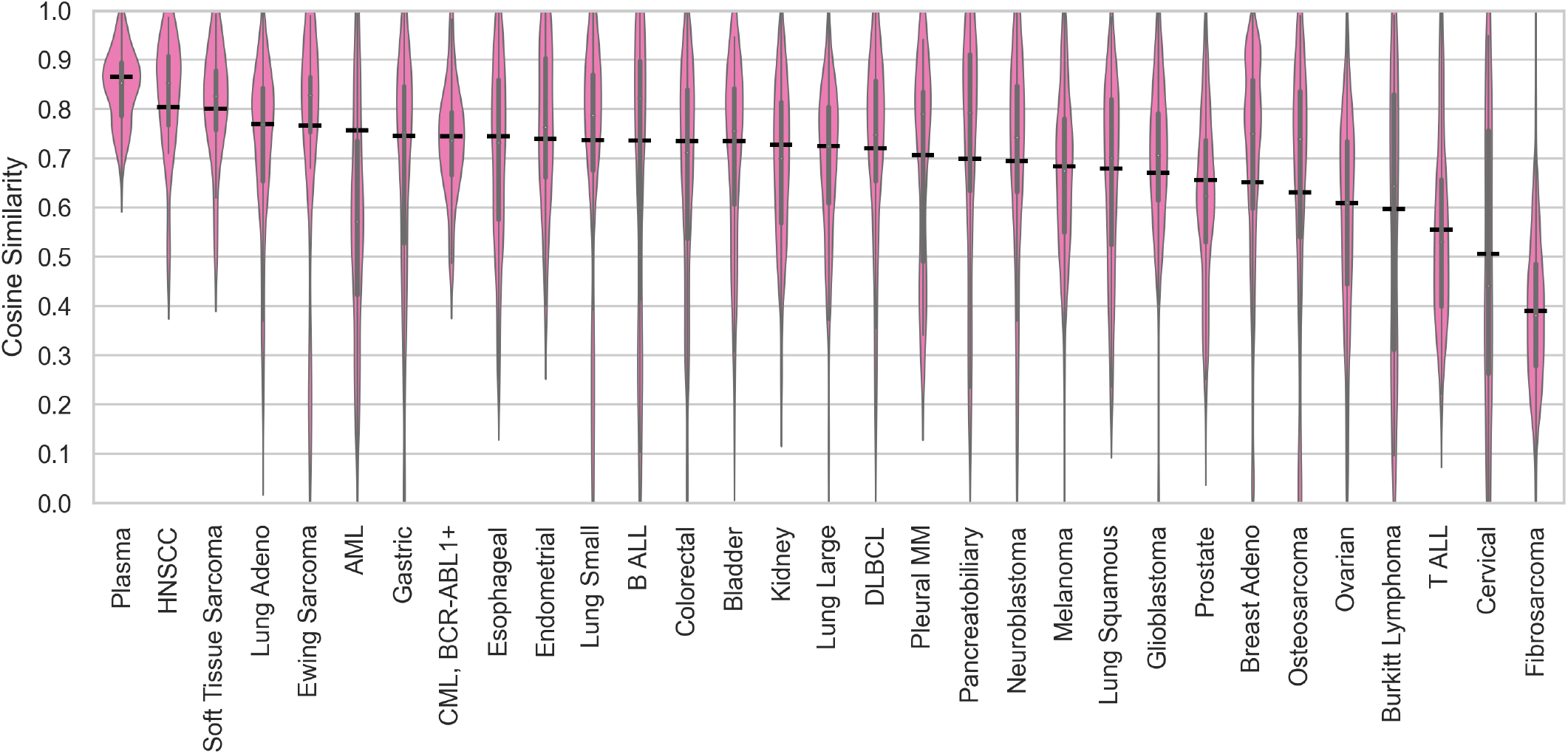
Pairwise similaritites of the same cell line instances per cancer types (*≥* 3 samples). Cell line averages are marked with a black dash. Abbreviations: AML - acute myeloid leukemia, B ALL - B acute lymphoblastic leukemia, CML - chronic myelogenous leukemia, DLBCL - diffuse large B-cell lymphoma, Plasma - plasma cell neoplasm, Pleural MM - pleural malignant mesothelioma, T ALL - T acute lymphoblastic leukemia

### Selection of representative cell lines

Our analysis, utilizing a SVM model trained on 31 distinct cancer types and complemented by visual assessment of CNV profiles, has yielded a shortlist of cell lines that serve as strong candidates for faithfully representing instances of 15 types of primary neoplasia (Table 1). Notably, these cell lines were consistently predicted across all 31 cancer types within the SVM model, suggesting a broader applicability rather than overtraining towards a specific subtype match. Out of all 31 cancer types analysed, we were able to determine candid models for 15 types. The highest number of accurate models were identified for breast adenocarcinomas where prediction accuracy was also the highest. We were also able to pinpoint 3 models for AML despite overall low prediction accuracy for this hematologic neoplasia.

**Table 1.** Cell lines with good correspondence to native cancer types.

| **Cancer Types** | **Cell Lines** |
| --- | --- |
| Acute Myeloid Leukemia | KO52 (CVCL_1321), KG-1 (CVCL_0374), MV4-11 (CVCL_0064) |
| Breast Adenocarcinoma | EFM-19 (CVCL_0253), ZR-75-1 (CVCL_0588), MDA-MB-134-VI (CVCL_0617), MDA-MB-361 (CVCL_0620), CAL-148 (CVCL_1106), CAL-85-1 (CVCL_1114), CAMA-1 (CVCL_1115), COLO 824 (CVCL_1136), DU4475 (CVCL_1183), Evsa-T (CVCL_1207), HCC1008 (CVCL_1244), HCC1143 (CVCL_1245), HCC1428 (CVCL_1252), HCC1500 (CVCL_1254), HCC1599 (CVCL_1256), HCC2218 (CVCL_1263), (CVCL_1267), HCC38 (CVCL_1270), MDA-MB-175-VII (CVCL_1400), MFM-223 (CVCL_1408), ZR-75-30 (CVCL_1661), UACC-812 (CVCL_1781), UACC-893 (CVCL_1782), EFM-192A (CVCL_1812), HDQ-P1 (CVCL_2067), JIMT-1 (CVCL_2077), KPL-1 (CVCL_2094), BT-483 (CVCL_2319), VP229 (CVCL_2754), VP267 (CVCL_2755), OCUB-F (CVCL_3352), HCC712 (CVCL_3378), SUM190PT (CVCL_3423), SUM44PE (CVCL_3424), MDA-MB-468GFP (CVCL_DH83) |
| Colorectal Carcinoma | COLO 320DM (CVCL_0219), SW403 (CVCL_0545), SW948 (CVCL_0632), LS1034 (CVCL_1382), NCI-H508 (CVCL_1564), SW1463 (CVCL_1718), SW626 (CVCL_1725), COLO 201 (CVCL_1987), COLO 206F (CVCL_1988), WiDr (CVCL_2760) |
| Diffuse Large B-Cell Lymphoma | HT (CVCL_1290), OCI-Ly10 (CVCL_8795), OCI-Ly19 (CVCL_1878), Ri-1 (CVCL_1885), SU-DHL-4 (CVCL_0539), SU-DHL-5 (CVCL_1735) |
| Esophageal Carcinoma | KYSE-140 (CVCL_1347), KYSE-150 (CVCL_1348), KYSE-270 (CVCL_1350), KYSE-30 (CVCL_1351), KYSE-510 (CVCL_1354), OE21 (CVCL_2661), OE33 (CVCL_0471) |
| Glioblastoma | DK-MG (CVCL_1173), GaMG (CVCL_1226), SF295 (CVCL_1690), SNB-75 (CVCL_1706), YKG-1 (CVCL_1796) |
| Head and Neck Squamous Cell Carcinoma | SCC-25 (CVCL_1682), BICR 22 (CVCL_2310), CAL-27 (CVCL_1107) |
| Kidney Carcinoma | 769-P (CVCL_1050), A-498 (CVCL_1056), A-704 (CVCL_1065), ACHN (CVCL_1067), CAL-54 (CVCL_1111), VMRC-RCW (CVCL_1790), VMRC-RCZ (CVCL_1791), UM-RC-6 (CVCL_2741), KMRC-1 (CVCL_2983), KMRC-2 (CVCL_2984), SLR26 (CVCL_V612) |
| Lung Adenocarcinoma | HCC1171 (CVCL_5126), HCC2450 (CVCL_5133), NCI-H1437 (CVCL_1472), NCI-H1693 (CVCL_1488), NCI-H1993 (CVCL_1512), NCI-H2030 (CVCL_1517), NCI-H2087 (CVCL_1524), NCI-H2291 (CVCL_1546), NCI-H650 (CVCL_1575), NCI-H820 (CVCL_1592), NCI-H920 (CVCL_1599), RERF-LC-KJ (CVCL_1654) |
| Lung Small Cell Carcinoma | COR-L279 (CVCL_1140) |
| Lung Squamous Cell Carcinoma | VMRC-LCP (CVCL_1788), LC-1/sq (CVCL_3008) |
| Melanoma | C32 (CVCL_1097), C32TG (CVCL_2324), COLO 741 (CVCL_1133), COLO 829 (CVCL_1137), G-361 (CVCL_1220), Hs 936.T (CVCL_1033), HT-144 (CVCL_0318), IGR-37 (CVCL_2075), Ma-Mel-36 (CVCL_A171), Malme-3M (CVCL_1438), SK-MEL-1 (CVCL_0068), SK-MEL-103 (CVCL_6069), SK-MEL-19 (CVCL_6025), SK-MEL-199 (CVCL_6104), SK-MEL-29 (CVCL_6031), SK-MEL-30 (CVCL_0039), UACC-257 (CVCL_1779), WM1193C (CVCL_C265), WM164 (CVCL_7928), WM1799 (CVCL_A341), WM266-4 (CVCL_2765), WM3060 (CVCL_6796), WM3066 (CVCL_C270), WM3208V (CVCL_L028), WM3248 (CVCL_6798), WM51 (CVCL_6995), WM852 (CVCL_6804), WM902B (CVCL_6807) |
| Ovarian Carcinoma | 59M (CVCL_2291), COV318 (CVCL_2419), FU-OV-1 (CVCL_2047), OVCAR-4 (CVCL_1627), PEO1 (CVCL_2686), PEO4 (CVCL_2690), PEO6 (CVCL_2691) |
| Pancreatobiliary Carcinoma | SU.86.86 (CVCL_3881), Capan-1 (CVCL_0237) |
| Prostate Carcinoma | NCI-H660 (CVCL_1576), VCaP (CVCL_2235) |

**Table 2.** Cancer types used in our analysis.

| Code | Label | CL | T |
| --- | --- | --- | --- |
| NCIT:C5214 | Breast Adenocarcinoma | 829 | 10756 |
| NCIT:C3224 | Melanoma | 689 | 1999 |
| NCIT:C3512 | Lung Adenocarcinoma | 472 | 4172 |
| NCIT:C2955 | Colorectal Carcinoma | 293 | 4787 |
| NCIT:C4908 | Ovarian Carcinoma | 280 | 2851 |
| NCIT:C4917 | Lung Small Cell Carcinoma | 212 | 318 |
| NCIT:C8851 | Diffuse Large B-Cell Lymphoma | 141 | 2797 |
| NCIT:C4911 | Gastric Carcinoma | 141 | 2278 |
| NCIT:C9384 | Kidney Carcinoma | 133 | 2569 |
| NCIT:C3058 | Glioblastoma | 115 | 3923 |
| NCIT:C34447 | Head and Neck Squamous Cell Carcinoma | 112 | 1848 |
| NCIT:C166418 | Pancreatobiliary Carcinoma | 107 | 1786 |
| NCIT:C3270 | Neuroblastoma | 107 | 1761 |
| NCIT:C3171 | Acute Myeloid Leukemia | 93 | 3805 |
| NCIT:C3513 | Esophageal Carcinoma | 87 | 1738 |
| NCIT:C3493 | Lung Squamous Cell Carcinoma | 80 | 1802 |
| NCIT:C4912 | Bladder Carcinoma | 71 | 1689 |
| NCIT:C4863 | Prostate Carcinoma | 67 | 3890 |
| NCIT:C4450 | Lung Large Cell Carcinoma | 65 | 92 |
| NCIT:C7376 | Pleural Malignant Mesothelioma | 56 | 211 |
| NCIT:C9039 | Cervical Carcinoma | 54 | 905 |
| NCIT:C2912 | Burkitt Lymphoma | 51 | 225 |
| NCIT:C3043 | Fibrosarcoma | 50 | 138 |
| NCIT:C4665 | Plasma Cell Neoplasm | 50 | 1617 |
| NCIT:C7558 | Endometrial Carcinoma | 48 | 819 |
| NCIT:C9145 | Osteosarcoma | 47 | 309 |
| NCIT:C4817 | Ewing Sarcoma | 43 | 182 |
| NCIT:C3183 | T Acute Lymphoblastic Leukemia | 40 | 238 |
| NCIT:C3174 | Chronic Myelogenous Leukemia, BCR-ABL1+ | 36 | 253 |
| NCIT:C8644 | B Acute Lymphoblastic Leukemia | 32 | 275 |
| NCIT:C9306 | Soft Tissue Sarcoma | 28 | 1196 |

To further verify the quality of our selection we calculated pairwise cosine similarities between the native cancer samples and instances of each individual cell lines (Supplementary Materials). These data indicate that for some cancer types, such as prostate carcinoma, suggested cell lines indeed corresponded to predominant sample types. In general, although similarity scores between the suggested cell lines varied widely the trained model showed an ability to successfully identify promising candidates even with low overall CNV pattern similarity.

## Discussion

We analyzed the molecular variability of over 500 cell lines (around 3,400 samples) using copy number variation profiles data. Overall, different instances of the same cell line displayed mostly limited variability. Due to the shared origin of these samples, this consistency in prediction is unsurprising. Higher similarity of different cancer cell lines of the same type to each other may be due to cell lines originating from a smaller clonal population and therefore being more stable. This trade-off presents a key challenge: While simplified models may offer stability and ease of analysis, they inherently struggle to capture the full spectrum of disease heterogeneity.

Studies with cancer cell lines have demonstrated that genetic drift occurs in cancer cell lines and significantly affects their pharmacogenetic properties (22, 23). Our results suggest that the heterogeneity of cancer cell lines is limited, especially when compared to native cancers. However, further analyses are required to determine the full scope of the genetic drift in the cell lines. Additionally, as apparent genetic drift in cancer cell lines can be the result of positive selection (22), some level of heterogeneity is inevitable.

The results of this study confirm that cancer cell lines indeed generally harbor a higher amount of CNVs compared to native cancers but also generally display CNVs shared with native cancers of the same diagnosis (Table 3, Fig. 8 and (9, 10)). The highest differences in CNV coverage were detected for leukemias, in particular for CML which can be attributed to the ease of using blast phase CML for generation of immortal cell lines (13) with inherent differences in genomic stability. Furthermore, this demonstrates that not all stages of cancers are well represented in *in vitro* models.

**Table 3.** Cosine similarities of CNV frequencies between Tumor and Cell Line.

| Cancer Type | Cosine Similarity |
| --- | --- |
| Lung Small Cell Carcinoma | 0.95 |
| Esophageal Carcinoma | 0.94 |
| Gastric Carcinoma | 0.94 |
| Lung Adenocarcinoma | 0.94 |
| Melanoma | 0.93 |
| Head and Neck Squamous Cell Carcinoma | 0.93 |
| Ovarian Carcinoma | 0.92 |
| Pancreatobiliary Carcinoma | 0.92 |
| Breast Adenocarcinoma | 0.92 |
| Colorectal Carcinoma | 0.90 |
| Kidney Carcinoma | 0.89 |
| Lung Squamous Cell Carcinoma | 0.89 |
| Pleural Malignant Mesothelioma | 0.88 |
| Glioblastoma | 0.87 |
| Cervical Carcinoma | 0.86 |
| Lung Large Cell Carcinoma | 0.86 |
| Bladder Carcinoma | 0.85 |
| Prostate Carcinoma | 0.84 |
| Endometrial Carcinoma | 0.83 |
| Diffuse Large B-Cell Lymphoma | 0.83 |
| Osteosarcoma | 0.82 |
| Plasma Cell Neoplasm | 0.81 |
| Burkitt Lymphoma | 0.80 |
| Ewing Sarcoma | 0.80 |
| Neuroblastoma | 0.80 |
| Acute Myeloid Leukemia | 0.78 |
| Soft Tissue Sarcoma | 0.73 |
| T Acute Lymphoblastic Leukemia | 0.70 |
| Chronic Myelogenous Leukemia, BCR-ABL1+ | 0.58 |
| Fibrosarcoma | 0.55 |
| B Acute Lymphoblastic Leukemia | 0.39 |

**Table 4.** Accuracies for the SVM model. Correct test % shows the percentage of correct predictions in the testing phase. Correct % shows the percentage of correct predictions of cell line diagnoses.

| Actual | Label | Correct Test % | Correct % |
| --- | --- | --- | --- |
| NCIT:C5214 | Breast Adenocarcinoma | 78.89 | 84.56 |
| NCIT:C3058 | Glioblastoma | 78.32 | 63.48 |
| NCIT:C2955 | Colorectal Carcinoma | 71.15 | 62.80 |
| NCIT:C3513 | Esophageal Carcinoma | 32.83 | 56.32 |
| NCIT:C9384 | Kidney Carcinoma | 72.22 | 54.89 |
| NCIT:C3224 | Melanoma | 60.11 | 53.70 |
| NCIT:C3270 | Neuroblastoma | 77.18 | 46.73 |
| NCIT:C8851 | Diffuse Large B-Cell Lymphoma | 55.28 | 46.10 |
| NCIT:C4665 | Plasma Cell Neoplasm | 68.00 | 44.00 |
| NCIT:C4863 | Prostate Carcinoma | 64.36 | 41.79 |
| NCIT:C3512 | Lung Adenocarcinoma | 62.00 | 37.08 |
| NCIT:C34447 | Head and Neck Squamous Cell Carcinoma | 37.75 | 36.61 |
| NCIT:C4908 | Ovarian Carcinoma | 56.56 | 35.36 |
| NCIT:C166418 | Pancreatobiliary Carcinoma | 34.38 | 28.04 |
| NCIT:C3171 | Acute Myeloid Leukemia | 90.51 | 19.35 |
| NCIT:C9039 | Cervical Carcinoma | 33.02 | 16.67 |
| NCIT:C4917 | Lung Small Cell Carcinoma | 40.51 | 15.09 |
| NCIT:C3493 | Lung Squamous Cell Carcinoma | 56.37 | 10.00 |
| NCIT:C4911 | Gastric Carcinoma | 30.90 | 7.09 |
| NCIT:C4912 | Bladder Carcinoma | 42.30 | 7.04 |
| NCIT:C7376 | Pleural Malignant Mesothelioma | 9.62 | 5.36 |
| NCIT:C9145 | Osteosarcoma | 17.65 | 4.26 |
| NCIT:C7558 | Endometrial Carcinoma | 18.72 | 4.17 |
| NCIT:C8644 | B Acute Lymphoblastic Leukemia | 40.58 | - |
| NCIT:C9306 | Soft Tissue Sarcoma | 36.33 | - |
| NCIT:C4817 | Ewing Sarcoma | 13.46 | - |
| NCIT:C2912 | Burkitt Lymphoma | 4.17 | - |
| NCIT:C3043 | Fibrosarcoma | 2.86 | - |

**Table 5.** Accuracies for RF model. Correct test % shows the percentage of correct predictions in the testing phase. Correct % shows the percentage of correct predictions of cell line diagnoses.

| Actual | Label | Correct Test % | Correct % |
| --- | --- | --- | --- |
| NCIT:C5214 | Breast Adenocarcinoma | 87.26 | 88.78 |
| NCIT:C2955 | Colorectal Carcinoma | 72.83 | 70.31 |
| NCIT:C3058 | Glioblastoma | 79.34 | 59.13 |
| NCIT:C8851 | Diffuse Large B-Cell Lymphoma | 60.29 | 48.23 |
| NCIT:C34447 | Head and Neck Squamous Cell Carcinoma | 37.40 | 41.07 |
| NCIT:C9384 | Kidney Carcinoma | 69.22 | 40.60 |
| NCIT:C3270 | Neuroblastoma | 79.92 | 39.25 |
| NCIT:C3224 | Melanoma | 41.73 | 33.38 |
| NCIT:C4908 | Ovarian Carcinoma | 48.65 | 32.86 |
| NCIT:C4863 | Prostate Carcinoma | 67.26 | 28.36 |
| NCIT:C3512 | Lung Adenocarcinoma | 62.91 | 27.33 |
| NCIT:C4665 | Plasma Cell Neoplasm | 71.46 | 22.00 |
| NCIT:C3513 | Esophageal Carcinoma | 27.08 | 21.84 |
| NCIT:C3183 | T Acute Lymphoblastic Leukemia | 42.25 | 20.00 |
| NCIT:C3493 | Lung Squamous Cell Carcinoma | 51.33 | 15.00 |
| NCIT:C9145 | Osteosarcoma | 6.06 | 14.89 |
| NCIT:C3171 | Acute Myeloid Leukemia | 84.51 | 13.98 |
| NCIT:C166418 | Pancreatobiliary Carcinoma | 34.21 | 13.08 |
| NCIT:C3174 | Chronic Myelogenous Leukemia, BCR-ABL1+ | 36.99 | 11.11 |
| NCIT:C4912 | Bladder Carcinoma | 28.16 | 8.45 |
| NCIT:C4911 | Gastric Carcinoma | 29.09 | 7.09 |
| NCIT:C9039 | Cervical Carcinoma | 24.70 | 5.56 |
| NCIT:C4917 | Lung Small Cell Carcinoma | 29.90 | 5.19 |
| NCIT:C7558 | Endometrial Carcinoma | 25.67 | 2.08 |
| NCIT:C8644 | B Acute Lymphoblastic Leukemia | 58.75 | - |
| NCIT:C9306 | Soft Tissue Sarcoma | 28.12 | - |
| NCIT:C2912 | Burkitt Lymphoma | 22.39 | - |
| NCIT:C4817 | Ewing Sarcoma | 13.33 | - |
| NCIT:C7376 | Pleural Malignant Mesothelioma | 9.23 | - |
| NCIT:C4450 | Lung Large Cell Carcinoma | 8.00 | - |

**Fig. 8.**
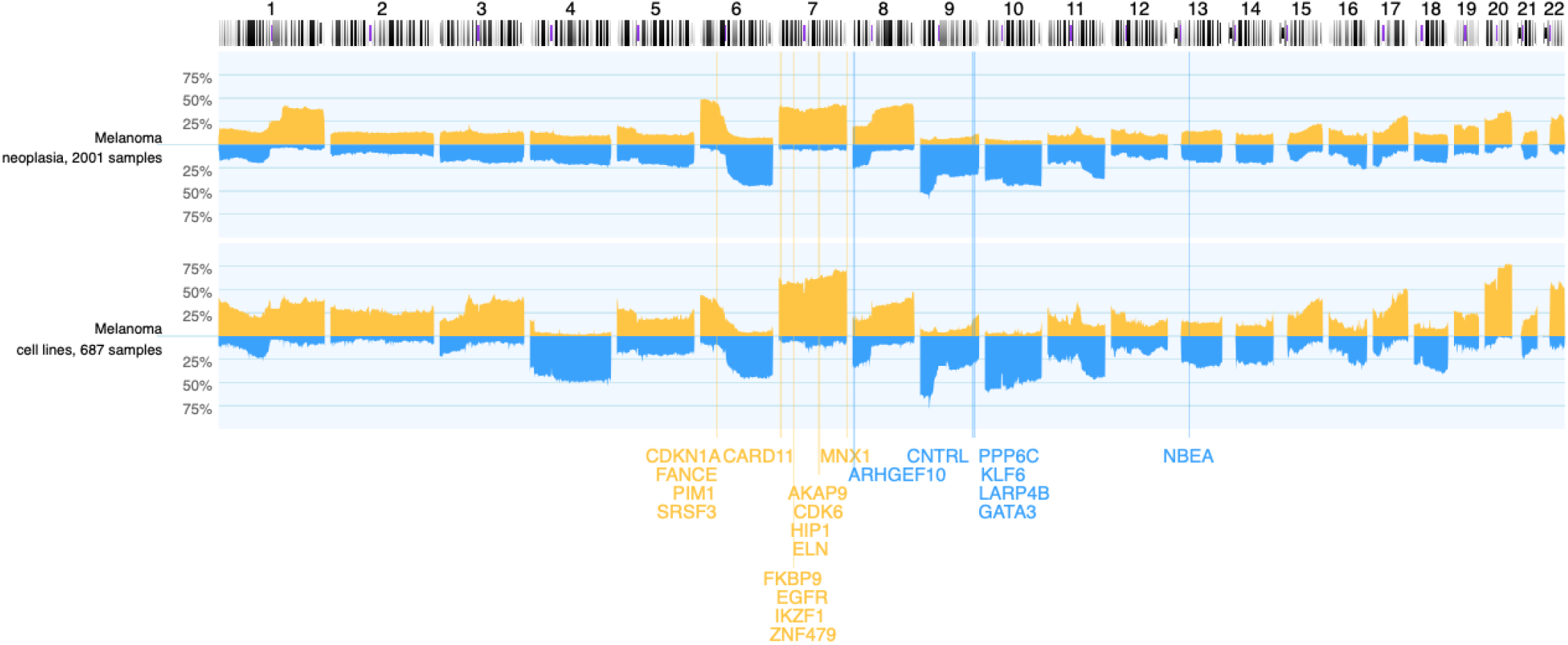
Frequency map of primary melanoma samples and melanoma cell lines annotated with genes in identified important features

We examined CNV samples of 31 different cancer types and showed that neoplasias exhibit high molecular variability (Fig. 3, Fig. 9). A useful strategy in taking advantage of cell lines would be to partition primary cancer into subsets and match these subsets to cell lines by similarity, a strategy we successfully employed to match cell lines to a subset of lung small cell carcinoma samples (Fig. 6). Overall, the current model may not fully account for the significant variations associated with within different cancer subtypes and potentially could be improved by a general approach including partitioning of cancer types and identification of cell lines that closely resemble those rather than unselected, potentially heterogeneous diagnostic classes.

**Fig. 9.**
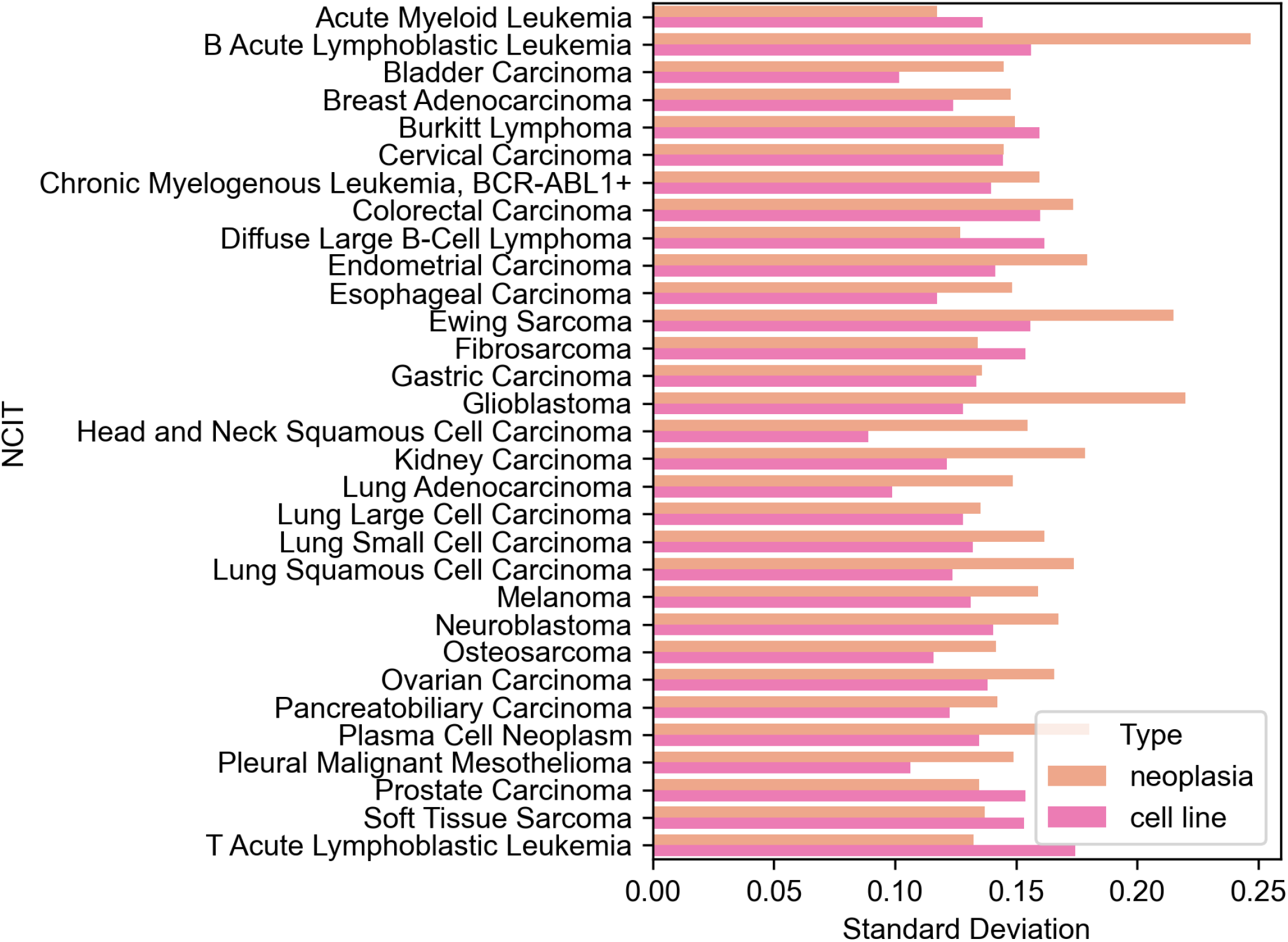
Standard deviations of cosine similarities between cancer cell lines and within neoplasia samples.

**Fig. 10.**
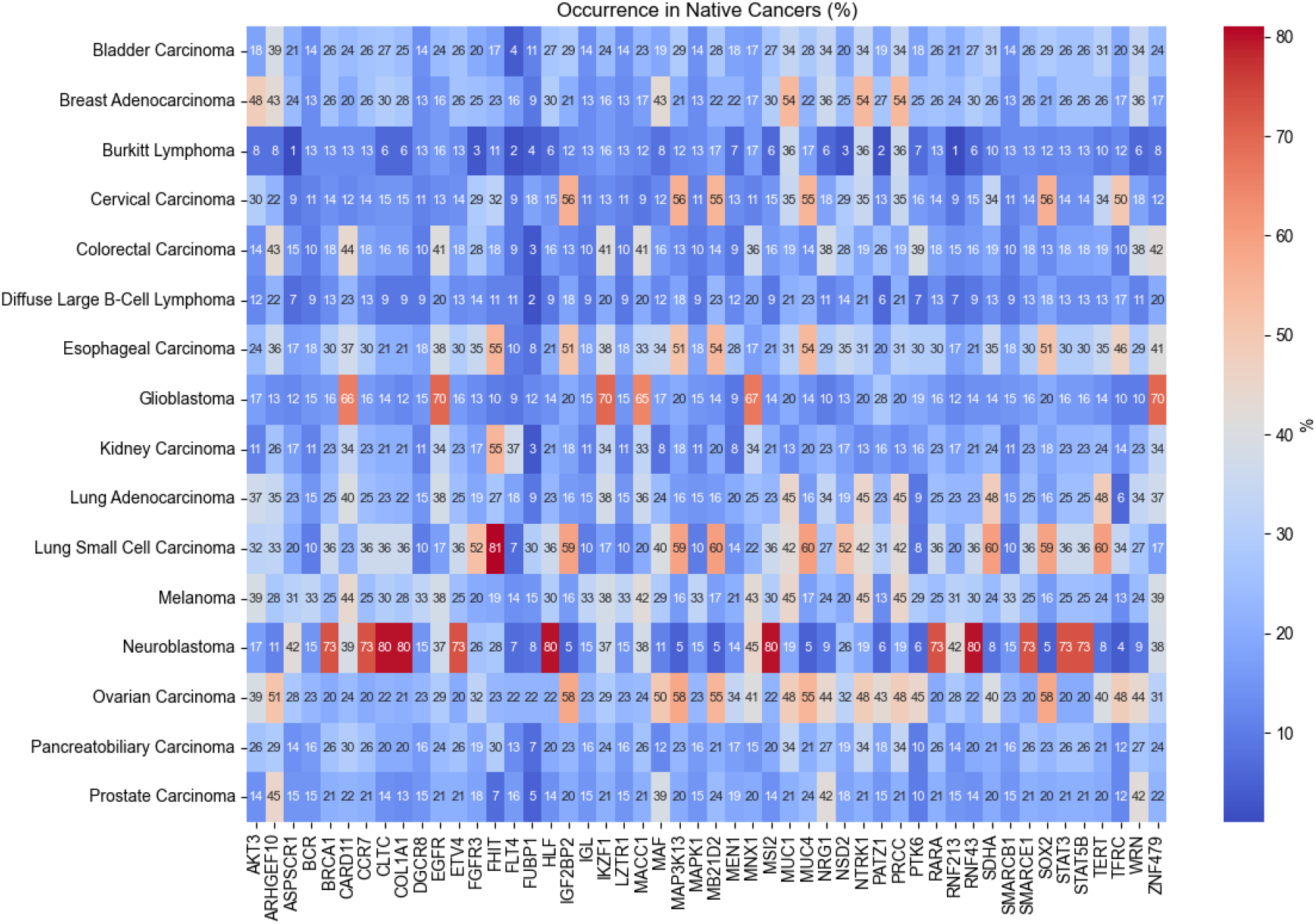
Percentage distribution of genes across native cancer samples of different types.

**Fig. 11.**
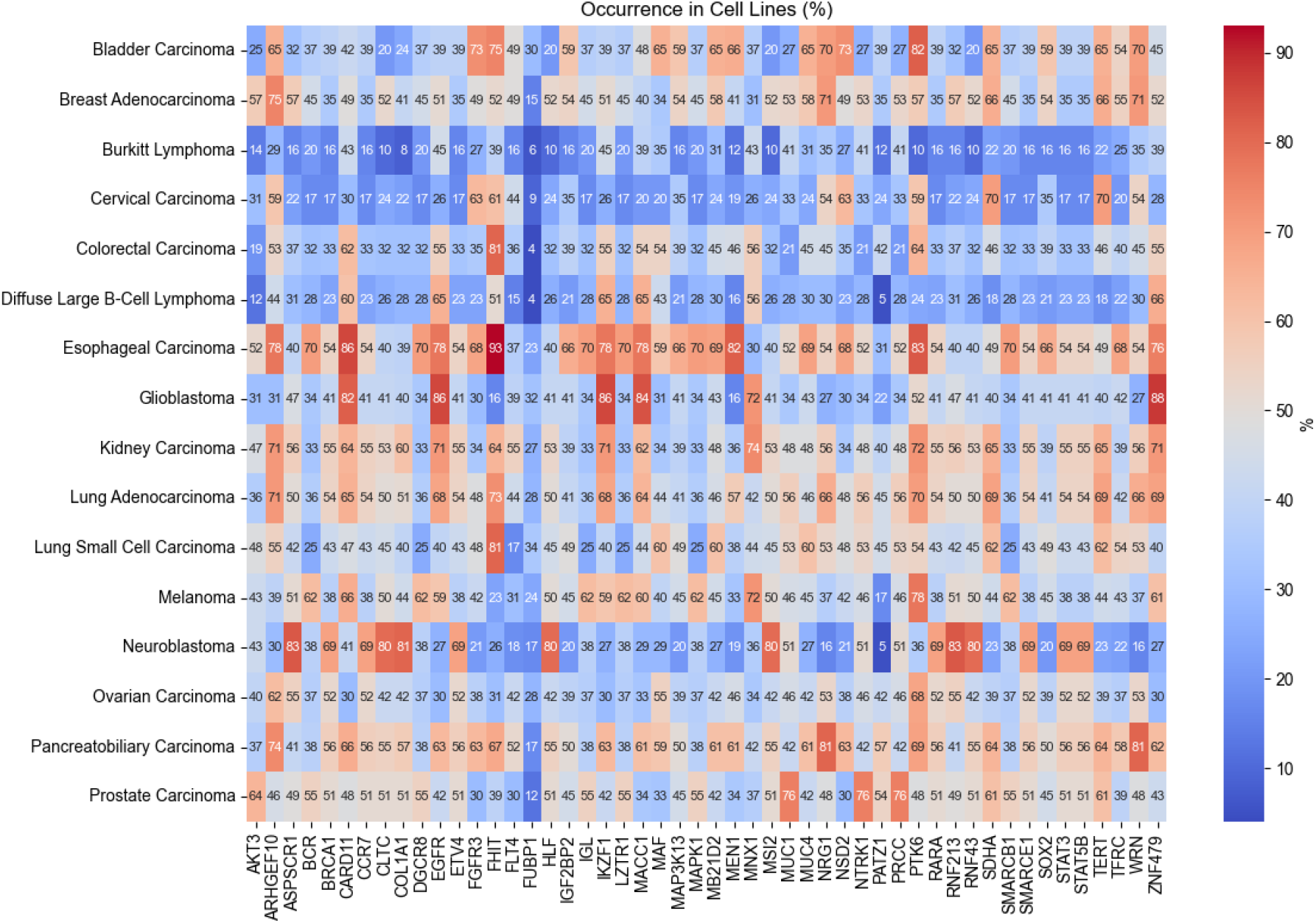
Percentage distribution of genes across cell line samples of different types.

We demonstrated that the prediction of cell line’s disease classification based on CNV patterns of native cancers was highly accurate for breast adenocarcinomas, colorectal carcinomas and glioblastomas. Using ML algorithms to classify cancers and cell lines also informs us about genomic features important for classifications. We showed that several well-known oncogenes and tumor suppressor genes might have influenced the decision-making process (Fig. 5). We also demonstrate that duplications are more important for class determination than deletions.

One effective approach to improving the performance of classification models on high-dimensional features like data from genomic screening experiments is to augment the sample size. For our particular analysis, addition of samples could lead to additional cell line matches while for underrepresented cancer types as well as a general approach consistent partitioning of cancer samples for subtype specific matching. Importantly our analyses hint at biases in cancer type representation through cell lines, at least such with accessible CNV profiling data.

In summary, despite the undeniable value of cancer cell lines in elucidating tumor biology and propelling advancements in precision medicine, the inherent genomic heterogeneity observed in cancer samples and across individuals needs to be accounted for. We provide a careful selection of the models corresponding best to the target disease to best capture the genomic intricacies of cancers. Data to create neoplasia subsets and match them to appropriate cell lines are available through the *Progenetix* and *cancercelllines*.*org* resources. Future steps in utilizing these resources could involve the creation of software tools to enable dynamic comparisons of cancer cell lines to native cancers.

## Supplementary Note 1: Supplementary materials

## Supplementary Note 2: Cosine similarities

Violin plots of cosine similarities between native cancer and cell line samples are included in this appendix. Average cosine similarities are marked by black lines on the violins. Cell lines suggested in the article are shown in red font. [pages=-]appendixB.pdf

## Notes

### Competing Interest Statement

The authors have declared no competing interest.

### Summary of Updates

Updated figures and figure descriptions. Details included to methods section.

